# PremPS: Predicting the Effects of Single Mutations on Protein Stability

**DOI:** 10.1101/2020.04.07.029074

**Authors:** Yuting Chen, Haoyu Lu, Ning Zhang, Zefeng Zhu, Shuqin Wang, Minghui Li

## Abstract

Protein stability is related to its functional activities, and effect on stability or misfolding could be one of the major disease-causing mechanisms of missense mutations. Here we developed a novel machine learning computational method PremPS, which predicts the effects of single mutations on protein stability by calculating the changes in unfolding Gibbs free energy. PremPS uses only ten evolutionary- and structure-based features and is parameterized on five thousand mutations. Our approach outperforms previous methods and shows a considerable improvement in estimating the effects of mutations increasing protein stability. In addition, PremPS presents an outstanding performance in predicting the pathogenicity of missense mutations using an experimental dataset composed of two thousand non-neutral and neutral mutations. PremPS can be applied to many tasks, including finding functionally important variants, revealing the molecular mechanisms of functional influences and protein design. It is freely available at https://lilab.jysw.suda.edu.cn/research/PremPS/.

**Key Points:**

- Considerable improvement in estimating the effects of mutations increasing protein stability;
- Comprehensive comparison with other 25 computational methods on different test sets;
- An outstanding performance in predicting the pathogenicity of missense mutations;
- PremPS employs only ten distinct features belonging to six categories, and the most important feature describes evolutionary conservation of the site;
- The webserver allows to do large-scale mutational scanning and takes about ten minutes to perform calculations for one thousand mutations from a normal size protein.

## INTRODUCTION

Protein stability is one of the most important factors that characterize protein function, activity and regulation (1). Missense mutations can lead to protein dysfunction by affecting their stabilities and interactions with other biological molecules (2-6). Several studies have shown that the mutations are deleterious due to decreasing/enhancing the stability of the corresponding protein (7-12). To quantify the effects on protein stability requires estimating the changes in folding/unfolding Gibbs free energy induced by mutations. Experimental measurements of protein stability changes are laborious and appropriate only for proteins that can be purified (13). Therefore, computational prediction is urgently required, which would help the prioritization of potentially functionally important variants and become vital to many fields, such as medical applications (14) and protein design (15).

A lot of computational approaches have been developed in the last decades to predict the effects of single mutations on protein stability (16-42). The vast majority of them are machine learning approaches and based on protein 3D structures. They are different in terms of algorithms used for building models, structural optimization procedures or features of energy functions. The prediction performances of these methods have been assessed and compared using several different datasets of experimentally characterized mutants (11,43-46). The results indicate that all methods showed a correct trend in the predictions but with inconsistent performances for different test sets. A majority of methods presented moderate or low accuracies when applied to the independent test sets, and PoPMuSiC (16), FoldX (23) and mCSM (17) showed relatively better performances in comparison with other methods.

The machine learning approaches are prone to have overfitting problems (47), namely their predictions tend to be biased towards the characteristics of learning datasets. The training data sets available so far with experimentally determined protein stability changes are enriched with destabilizing mutations (16,48). Thus, the vast majority of predictors that did not consider the unbalance of training dataset showed a better performance for predicting destabilizing than stabilizing mutations (49,50). A recent study constructed a balanced data set with an equal number of destabilizing and stabilizing mutations and was used to assess the performance of 14 methods (51). The results showed that the 14 widely used predictors present a strong bias toward predicting the destabilizing mutations. Correcting for such bias in method’s performance is not a trivial task, which requires enriching the training set with stabilizing mutations and developing new energy functions.

To address this issue, we developed PremPS that uses a novel scoring function composed of only ten features and trains on a balanced dataset including five thousand mutations, half of which belong to destabilizing mutations and the remaining half are stabilizing mutations. PremPS has been validated to perform significantly better than other methods, especially in predicting the effects of stabilizing mutations. PremPS can be applied to many tasks, including finding functionally important variants, revealing the molecular mechanisms of functional influences and protein design.

## MATERIALS AND METHODS

### Experimental dataset for parameterizing PremPS

ProTherm database is a collection of thermodynamic parameters for wild-type and mutant proteins (48). It contains unfolding Gibbs free energy changes that provide important clues for estimating and interpreting the relationship among structure, stability and function of proteins and their mutants. It is frequently used as training template for developing *in silico* prediction approaches.

S2648 dataset includes 2648 single-point mutations from 131 globular proteins (Figure S1), which was derived from the ProTherm database and compiled by (16). It was used as the training dataset of PoPMuSiC (16), mCSM (17) and DUET (18) methods. Here, we also used the mutations and their unfolding Gibbs free energy changes from the S2648 to parameterize PremPS model. Then, we applied the following criteria to update the protein 3D structures: structure obtained/extracted from monomer or homomer is preferred over heteromer; wild-type protein structure is preferred over mutant; structure with minimal number of ligands is used; crystal structure is preferred over NMR, and higher resolution structure is chosen. The multimeric state of each protein was either assigned by manually checking the corresponding references used to measure protein stability changes or retrieved from the PQS server (52).

Unfolding free energy change (ΔΔ*G*) of a system can be characterized as a state function where the ΔΔ*G*_*wt*→*mut*_ value of a forward mutation minus ΔΔ*G*_*mut*→*wt*_ of its reverse mutation should be equal to zero. Given the unbalanced nature of the S2648 dataset with 2080 destabilizing (decreasing stability, ΔΔ*G*_*exp*_ ≥ 0) and 568 stabilizing (increasing stability, ΔΔ*G*_*exp*_ < 0) mutations, we modelled their reverse mutations in order to establish a more accurate computational method. Therefore, the final training set for parameterizing PremPS model includes 5296 single mutations (it will be referred as S5296) (Table S1). The dataset can be downloaded from https://lilab.jysw.suda.edu.cn/research/PremPS/download/.

For the forward mutations (ΔΔ*G*_*wt*→*mut*_), 3D structures of wild-type proteins were obtained from the Protein Data Bank (PDB) (53). For the reverse mutations (ΔΔ*G*_*wt*→*mut*_), the 3D structures of mutants were produced by BuildModel module of FoldX (23) using wild-type protein structures as the templates.

### Experimental datasets used for testing

First, we used the following six datasets to assess and compare the predictive performance of PremPS and other 25 computational methods (16-40).

- S350, it is a randomly selected subset from S2648 including 350 mutations from 67 proteins (16). This dataset is widely used to compare the performance of different methods. During the comparison, all methods were retrained after removing S350 from their training sets.
- S605, it was compiled from Protherm database by (21) and contains 605 mutations from 58 proteins, which is the training dataset of Meta-predictor method (21).
- S1925, it includes 1925 mutations from 55 proteins evenly distributed over four major SCOP structural classes, which is the training dataset of AUTOMUTE method (24).
- S^sym^, a dataset was manually curated by (51). It contains 684 mutations, half of which belong to *forward mutations*, and the remaining half are *reverse mutations* with crystal structures of the corresponding mutant proteins available.
- S134, it consists of experimentally determined stability changes for 134 mutations from sperm-whale myoglobin (43), and six different high-resolution crystal structures of myoglobin were used for the energy calculation.
- p53, it includes 42 mutations within the DNA binding domain of the protein p53 with experimentally determined thermodynamic effects, and the data was obtained from (17).

The number of mutations in each test set is shown in Table S1a, and the number of overlapped mutations between our training set of S5296 and each test set is presented in Table S1b.

Next, we removed the redundant mutations from the above datasets and the overlapped mutations with S5296, then established a combined independent test set. For the proteins with multiple 3D structures, we used the same criteria used in processing the training dataset of S2648 to choose the optimal structure. For the conflicting entries with multiple experimental measurements, if the difference between the maximal and minimal ΔΔ*G*_*exp*_ for this mutation is less than 1.0 kcal mol^-1^, we used the average value, otherwise we removed all entries (Figure S2). As a result, the combined independent test set contains 921 single mutations (it will be referred as S921) (Table S1a).

### The model of PremPS

The random forest (RF) regression scoring function of PremPS is composed of ten distinct features belonging to six categories (described below) and parameterized on S5296 dataset. The contribution of each category of features is shown in Table S2.

- *PSSM* score is the Position-Specific Scoring Matrix created by PSI-BLAST. It finds similar protein sequences for the query sequence in which the mutation occurs by searching all protein sequences in NCBI non-redundant database, then builds a PSSM from the resulting alignment (54). The default parameters were applied to construct PSSM profile.
- *ΔCS* represents the change of conservation after mutation calculated by PROVEAN method (55). The features of *PSSM* and *ΔCS* illustrate that the evolutionarily conserved sites may play an important role in protein folding.
- *ΔOMH* is the difference of hydrophobicity scale between mutant and wild-type residue type. The hydrophobicity scale (OMH) for each type of amino acid residue, obtained from the study of (56), was derived by considering the observed frequency of amino acid replacements among thousands of related structures.
- *SASA*_*pro*_ and *SASA*_*sol*_ is the solvent accessible surface area (SASA) of the mutated residue in the protein and in the extended tripeptide respectively. The SASA of a residue in the protein and in the extended tripeptide was calculated by DSSP program (57) and obtained from (58), respectively.
- *P*_*FWY*_, *P*_*RKDE*_ and *P*_*L*_ is the fraction of aromatic residues (F, W or Y), charged residues (R, K, D or E) and leucine (L) buried in the protein core, respectively. For instance, 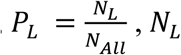 is the number of all leucine residues buried in the protein core and *N*_*All*_ is the total number of residues. If the ratio of solvent accessible surface area of a residue in the protein and in the extended tripeptide is less than 0.2 (59), we defined this residue as buried in the core of protein.
- *N*_*Hydro*_ and *N*_*Charg*_ is the number of hydrophobic (V, I, L, F, M, W, Y or C) and charged amino acids (R, K, D or E) at 23 sites in the protein sequence, the center of which is at the mutated site, respectively (60).

In addition to Random Forest, we also tried two other popular learning algorithms of Support Vector Machine (SVM) and eXtreme Gradient Boosting (XGBoost), and the results indicate that random forest regression model presents the best performance (Table S3).

PremPS takes about ten minutes to perform calculations for one thousand mutations from a normal size protein (∼ 300 residues). The running time for a single mutation per normal size protein is about four minutes, and it requires ∼ 0.4 second for each additional mutation.

### Cross-validation procedures

We performed four types of cross validation (CV1-CV4). For CV1 and CV2, we randomly chose 80% and 50% of mutations from the S5296 set respectively to train the model and used the remaining mutations for blind testing; the procedures were repeated 100 times. The number of mutations is not uniformly distributed over proteins (Figure S1), in order to conquer the bias toward the proteins with large number of mutations, we carried out the third type of cross validation (CV3). Namely, a subset was created by randomly sampling up to 20 mutations for each protein from S5296; the procedure was repeated 10 times and resulted in 1704 mutations in each subset. Then 80% mutations were randomly selected from each subset to train the model and the rest of mutations were used for testing, repeated 10 times. Next, we performed leave-one-protein-out validation (CV4), in which the model was trained on all mutations from 130 protein structures and the rest of protein/mutations were used to evaluate the performance. This procedure was repeated for each protein and its mutations. In all four described cross-validation procedures, during the training/test splits, the forward and their corresponding reverse mutations were retained in the same set, either training or testing.

### Statistical analysis and evaluation of performance

We used two measures of Pearson correlation coefficient (R) and root-mean-square error (RMSE) to verify the agreement between experimental and predicted values of unfolding free energy changes. All correlation coefficients reported in the paper are significantly different from zero with p-value smaller than 0.01 (t-test). RMSE (kcal mol^-1^) is the standard deviation of the residuals (prediction errors). For checking whether the difference in performance between PremPS and other methods is significant, we used Hittner2003 (61) and Fisher1925 (62) tests implemented in package *cocor* from R (63) to compare two correlation coefficients. Receiver operating characteristics (ROC) curves were compared with Delong tests (64).

To quantify the performance of different methods, we performed Receiver Operating Characteristics (ROC) and precision-recall analyses. True positive rate is defined as TPR=TP/(TP + FN) and false positive rate is defined as FPR=FP/(FP+TN) (TP: true positive; TN: true negative; FP: false positive; FN: false negative). In addition, Matthews correlation coefficient is calculated for estimating the quality of binary classification and accounting for imbalances in the labeled dataset:

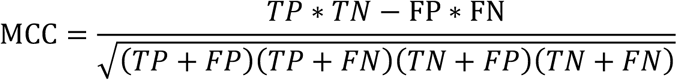

To compare across methods, the maximal MCC value was reported for each method through calculating the MCC across a range of thresholds.

## RESULTS and DISCUSSION

PremPS employs only ten features belonging to six categories and is constructed by random forest regression algorithm implemented in R randomForest package (65). The number of trees “ntree” is set to 500 and the number of features, randomly sampled as candidates for splitting at each node, “mtry” value is set to 3. All features have significant contribution to the model (Table S2). The performance of PremPS trained and tested on S5296 is shown in Figure 1a and Table S4. Pearson correlation coefficient between experimental and calculated unfolding free energy changes is 0.82 and the corresponding root-mean-square error and slope is 1.03 kcal mol^-1^ and 1.08 respectively.

**Figure 1.**
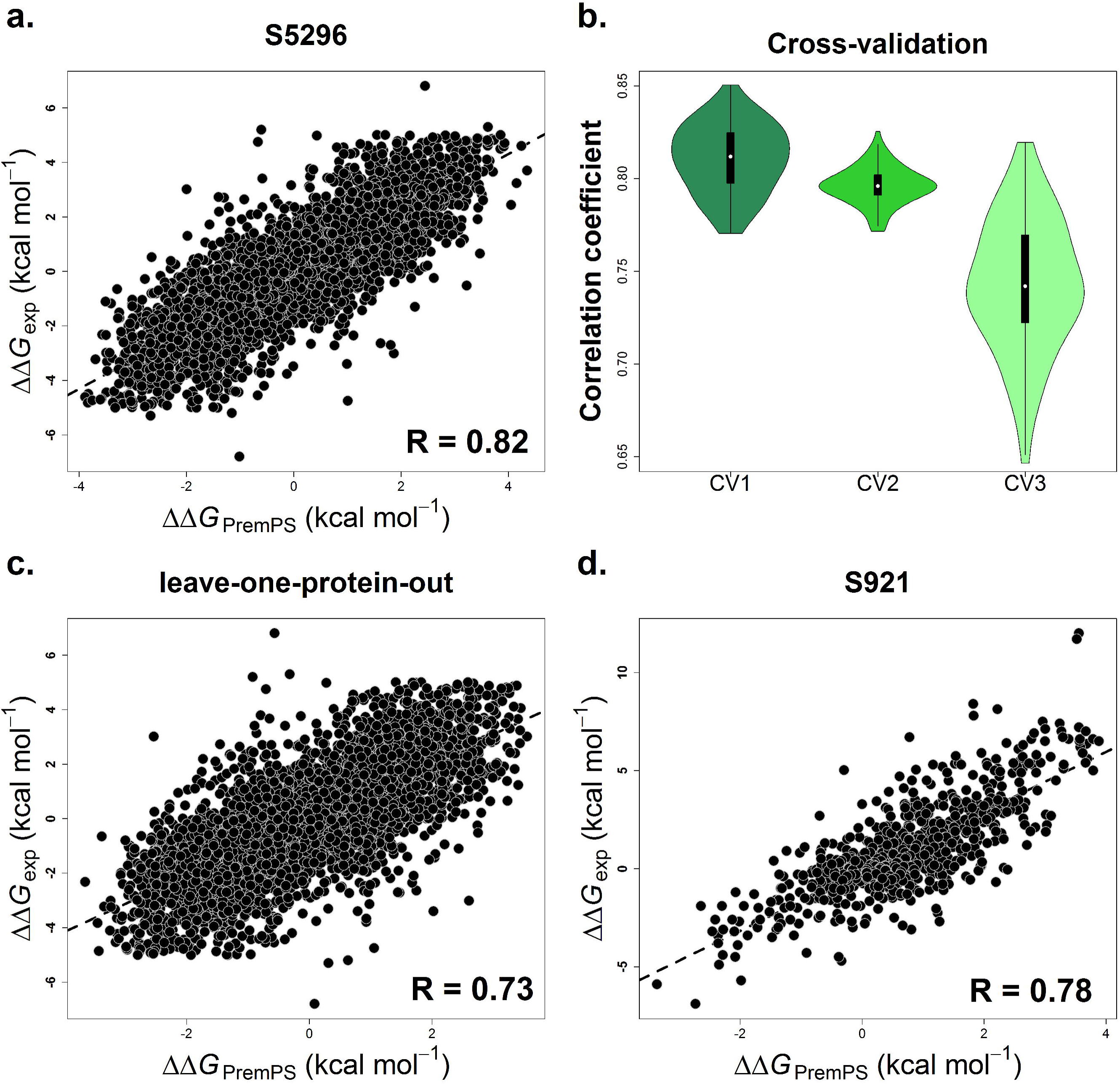
Pearson correlation coefficients between experimentally-determined and calculated values of changes in protein stability (ΔΔ*G*) for PremPS trained and tested on S5296 (a), applying three types of cross-validation (b) and leave-one-protein-out validation on S5296 (c), and tested on the independent dataset of S921 (d). See also Table S1 and Table S4.

### Performance on four types of cross-validation

Overfitting is one of the major concerns in machine learning, which may occur when the parameters are over-tuned to minimize the mean square deviations of predicted from experimental values in the training set. To overcome this problem, we performed four types of cross-validation (details were explained in Methods section), which is capable of estimating the performance of a method on previously unseen data. As shown in Figure 1b and Table S4, the correlation coefficient of each round in either CV1 or CV2 is higher than 0.77, and the mean values of R and RMSE for both cross-validation are ∼ 0.80 and ∼1.08 kcal mol^-1^ respectively across the 100 rounds. Taking the bias that the distribution of the number of mutations over proteins is not uniform into account, the CV3 cross-validation was performed. The mean values of R and RMSE is 0.74 and 1.21 kcal mol^-1^ respectively for CV3. Moreover, we evaluated the performance of PremPS on a low redundant set of proteins using leave-one-protein-out validation (CV4), the Pearson correlation coefficient of which reaches to 0.73 (Figure 1c) and RMSE = 1.23 kcal mol^-1^ (Table S4).

### Validation on independent test sets

First, we used six datasets of S350, S605, S1925, S^sym^, S134 and *p*53 (details about each set were shown in Methods section) to assess and compare the predictive performance of PremPS and other 25 computational methods that predict protein stability changes upon single mutations. The Pearson correlation coefficient and root-mean-square error for different methods applied on different datasets are presented in Table S5. The results indicate that the PremPS has the best performance, and the next are PoPMuSiC, FoldX and mCSM methods (the detailed descriptions are illustrated in the Table S5). Moreover, we removed the redundant mutations from the above six datasets and the overlapped mutations with the training set of S5296, and established the independent test set of S921. Then, we applied PremPS and other three methods of PoPMuSiC, FoldX and mCSM on this test set. The results reported in Figure 1d, Table 1 and Figure S3 demonstrate that PremPS achieves a high prediction accuracy with R of up to 0.78 and RMSE of 1.48 kcal mol^-1^, outperforming all three other methods.

**Table 1.** Comparison of methods’ performance on the independent test set of S921.

| Method | All mutations |  | Destabilizing |  | Stabilizing |  |
| --- | --- | --- | --- | --- | --- | --- |
|  | R | RMSE | R | RMSE | R | RMSE |
| PremPS | 0.78 | 1.48 | 0.72 | 1.54 | 0.60 | 1.33 |
| PoPMuSiC | 0.64 | 1.68 | 0.68<sup>#</sup> | 1.48 | - | 2.06 |
| FoldX | 0.57 | 2.06 | 0.56 | 1.99 | 0.22 | 2.21 |
| mCSM | 0.52 | 1.85 | 0.57 | 1.63 | - | 2.25 |

One of our goals is to improve the prediction accuracy for mutations increasing stability (stabilizing mutations). Table 1 and Table S4 show the considerable improvement for PremPS estimating the effects of stabilizing mutations. The R is 0.60 for PremPS tested on the stabilizing mutations from S921 set (Table 1). In addition, we evaluated the performance of PremPS when it was trained only on the forward mutation dataset of S2648 and tested on the S921. The correlation coefficients are 0.73 and 0.29 for destabilizing and stabilizing mutations, respectively (Table S6). The results confirm that the use of reverse mutations improved the performance of our model in estimating the effects of stabilizing mutations without compromising the prediction accuracy for destabilizing mutations.

### Performance on identification of highly destabilizing and stabilizing mutations

We carried out the ROC analysis in order to quantify the performance of PremPS in distinguishing highly destabilizing (ΔΔ*G* ≥ 1.0 kcal mol^-1^) and highly stabilizing (ΔΔ*G* ≤ -1.0 kcal mol^-1^) mutations from the others. Figure 2a and Figure S4 show excellent performance of PremPS in evaluating highly destabilizing/stabilizing mutations. We further subdivided mutations into two categories of core and surface according to the location of mutated site in the protein 3D structure (The definition was illustrated in the Methods section). We found that PremPS performed well for both categories and outperforms other methods on the S921 set (Figure 2b, Figure S5 and Table S6). The experimental ΔΔ*G* values for the majority of surface mutations are distributed near zero (Figure S5a), which might be the reason for the relatively lower correlation coefficient and RMSE compared to mutations in the core (RMSE is 1.81 and 1.11 kcal mol^-1^ for core and surface mutations, receptively) (Table S6).

**Figure 2.**
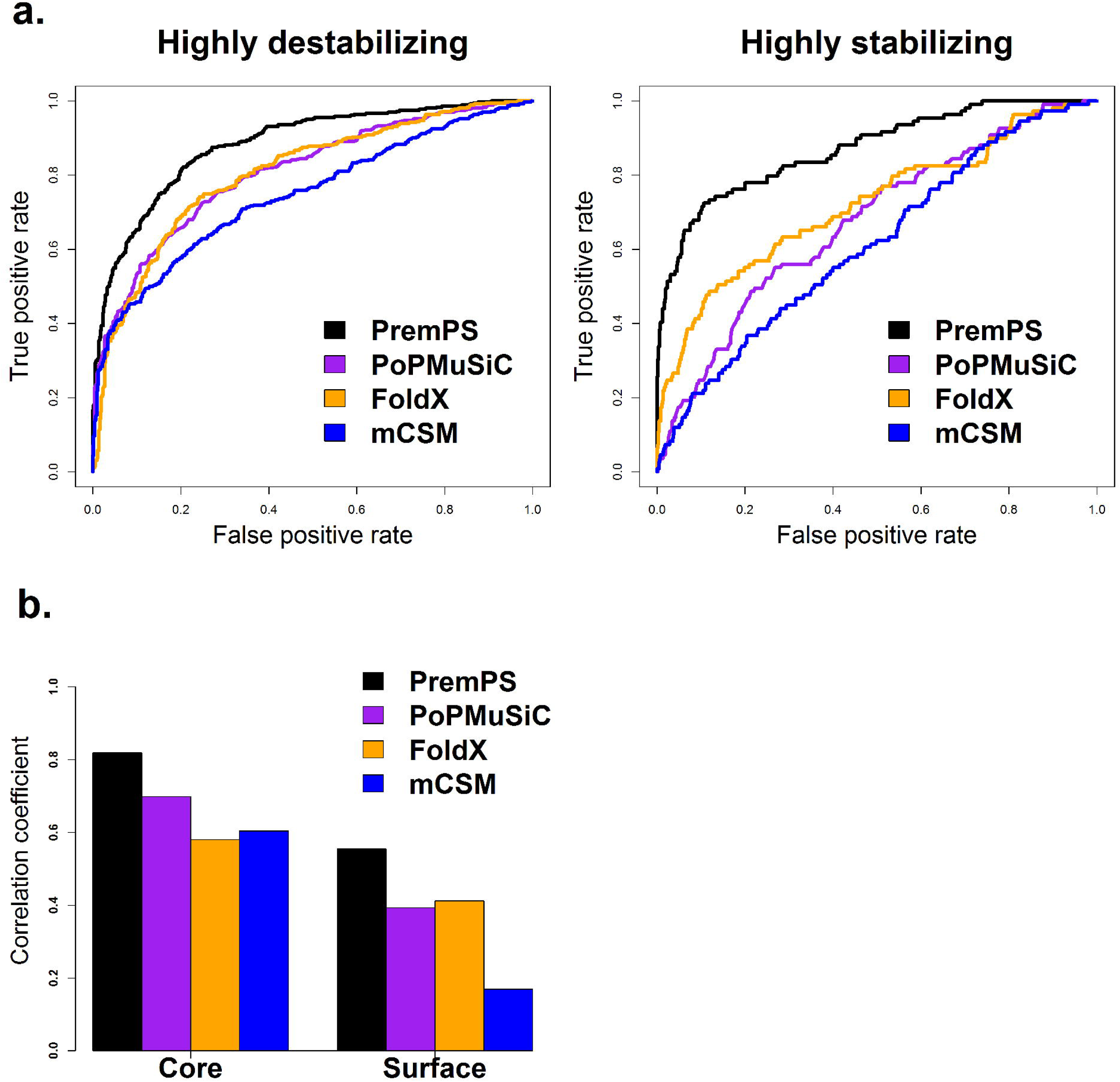
Comparative performance of PremPS and three other methods of PoPMuSiC, FoldX and mCSM on the independent test set of S921. (a) ROC curves for predicting highly destabilizing (ΔΔ*G*_*exp*_ ≥ 1 kcal mol^-1^) and highly stabilizing mutations (ΔΔ*G*_*exp*_ ≤ -1 kcal mol^-1^). PremPS has substantially higher AUC-ROC than other methods (p-value << 0.01, DeLong test, Figure S4b). (b) Pearson correlation coefficients between predicted and experimental ΔΔ*G* for mutations located in protein core and surface. The difference in R between PremPS and other methods is significant (p-value < 0.01, Hittner2003). More details are shown in the Figure S4 and S5.

### Performance on identification of functionally important mutations

Protein stability is related to its functional activity, and effect on stability could be one of the major disease-causing mechanisms of missense mutations. Therefore, we further checked the capability of PremPS on the identification of functionally important mutations. A previously proposed experimental dataset including 1139 non-neutral/deleterious and 4137 neutral/benign mutations was used here to evaluate the performance of PremPS in distinguishing non-neutral from neutral mutations (66). Among them 1037 non-neutral and 1159 neutral mutations (named as F2196 dataset) could be mapped to the corresponding protein 3D structures and allowed to calculate the unfolding free energy changes (Figure S6). We compared the performance of PremPS with 28 state-of-the-art computational methods developed for predicting the pathogenicity of missense mutations (67,68), such as CHASMplus (69), VEST4 (70), REVEL (71) and B-Score (66) (Figure S6). Table 2 and Figure S6 show that PremPS is capable of distinguishing non-neutral from neutral mutations. Since one protein could be mapped to several PDB structures, the number of mapped 3D structures for 50 proteins in F2196 dataset is 1554 (Fig. S6a). If the average value of protein stability changes calculated using all mapped structures was used, the performance of PremPS ranks in the middle (Fig. S6b), and if the maximum predicted absolute value was used for each non-neutral mutation and minimum predicted absolute value was used for each neutral mutation, PremPS performs better than all other methods (Table 2 and Fig. S6c).

**Table 2.**
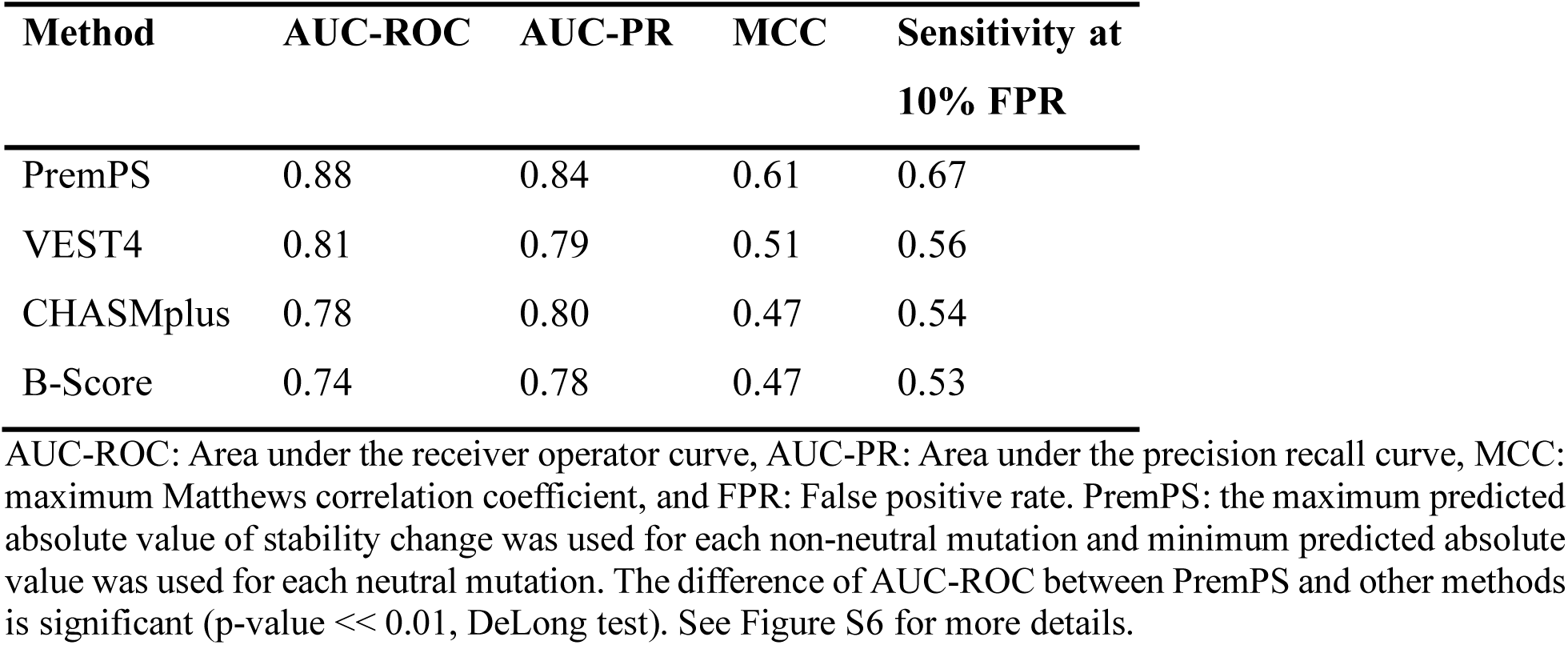
Comparison of the performance of different computational methods in distinguishing non-neutral from neutral mutations using the experimental dataset of F2196.

| **Method** | **AUC-ROC** | **AUC-PR** | **MCC** | **Sensitivity at**<br>**10% FPR** |
| --- | --- | --- | --- | --- |
| PremPS | 0.88 | 0.84 | 0.61 | 0.67 |
| VEST4 | 0.81 | 0.79 | 0.51 | 0.56 |
| CHASMplus | 0.78 | 0.80 | 0.47 | 0.54 |
| B-Score | 0.74 | 0.78 | 0.47 | 0.53 |

AUC-ROC: Area under the receiver operator curve, AUC-PR: Area under the precision recall curve, MCC: maximum Matthews correlation coefficient, and FPR: False positive rate. PremPS: the maximum predicted absolute value of stability change was used for each non-neutral mutation and minimum predicted absolute value was used for each neutral mutation. The difference of AUC-ROC between PremPS and other methods is significant (p-value $\ll 0.01$ , DeLong test). See Figure S6 for more details.

## ONLINE WEBSERVER

### Input

The 3D structure of a protein is required by the webserver, and the user can provide the Protein Data Bank (PDB) code or upload the coordinate file. When the user provides the PDB code, biological assemblies or asymmetric unit can be retrieved from the Protein Data Bank (Fig. 3). After the structure is retrieved correctly, the server will display a 3D view colored by protein chains and list the corresponding protein name (Fig. S7). At the second step one or multiple chains that must belong to one protein can be assigned to the following energy calculation. The third step is to select mutations and three options are provided: “Upload Mutation List”, “Alanine Scanning for Each Chain” and “Specify One or More Mutations Manually” (Fig. 3 and Fig. S7). “Upload Mutation List” allows users to upload a list of mutations for large-scale mutational scans. “Alanine Scanning for Each Chain” allows users to perform alanine scanning for each chain. In the option of “Specify One or More Mutations Manually”, users can not only perform calculations for specified mutations but also allow to view the mutated residues in the protein structure.

**Figure 3.**
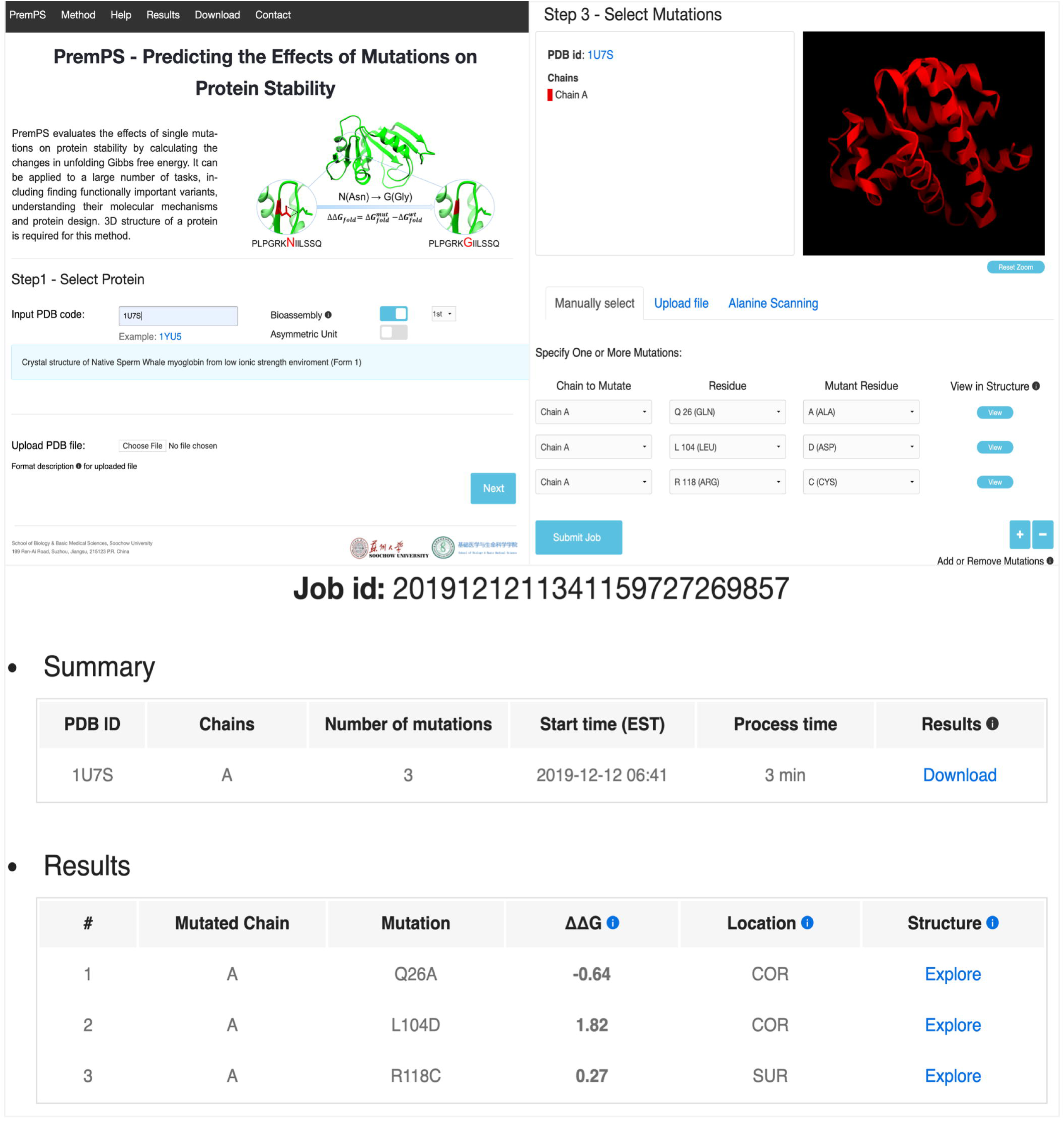
Left corner: the entry page of PremPS server; right corner: the third step for selecting mutations and three options are provided: “Specify One or More Mutations Manually”, “Upload Mutation List” and “Alanine Scanning for Each Chain”, see also Figure S7; and bottom: final results, see also Figure S8.

### Output

For each mutation of a protein, the PremPS server provides the following results (Figure 3): ΔΔ*G* (kcal mol^-1^), predicted unfolding free energy change induced by a single mutation (positive and negative sign corresponds to destabilizing and stabilizing mutations, respectively); location of the mutation occurred (COR: core or SUR: surface), a residue is defined as buried in the protein core if the ratio of solvent accessible surface area of this residue in the protein and in the extended tripeptide is less than 0.2, otherwise it is located on the surface of protein. In addition, for each mutation, the PremPS outputs the contribution of each feature in the target function and provides an interactive 3D viewer showing the non-covalent interactions between the mutated site and its adjacent residues, generated by Arpeggio (72) (Fig. S8).

## ACKNOWLEDGEMENT

This research was supported by the National Natural Science Foundation of China (Grant No. 31701136), Natural Science Foundation of Jiangsu Province, China (Grant No. BK20170335), and the Priority Academic Program Development of Jiangsu Higher Education Institutions.

## Competing interests

The authors declare no competing interests.

