## Supplementary Figures and Tables for "PremPS: Predicting the Effects of Single Mutations on Protein Stability"

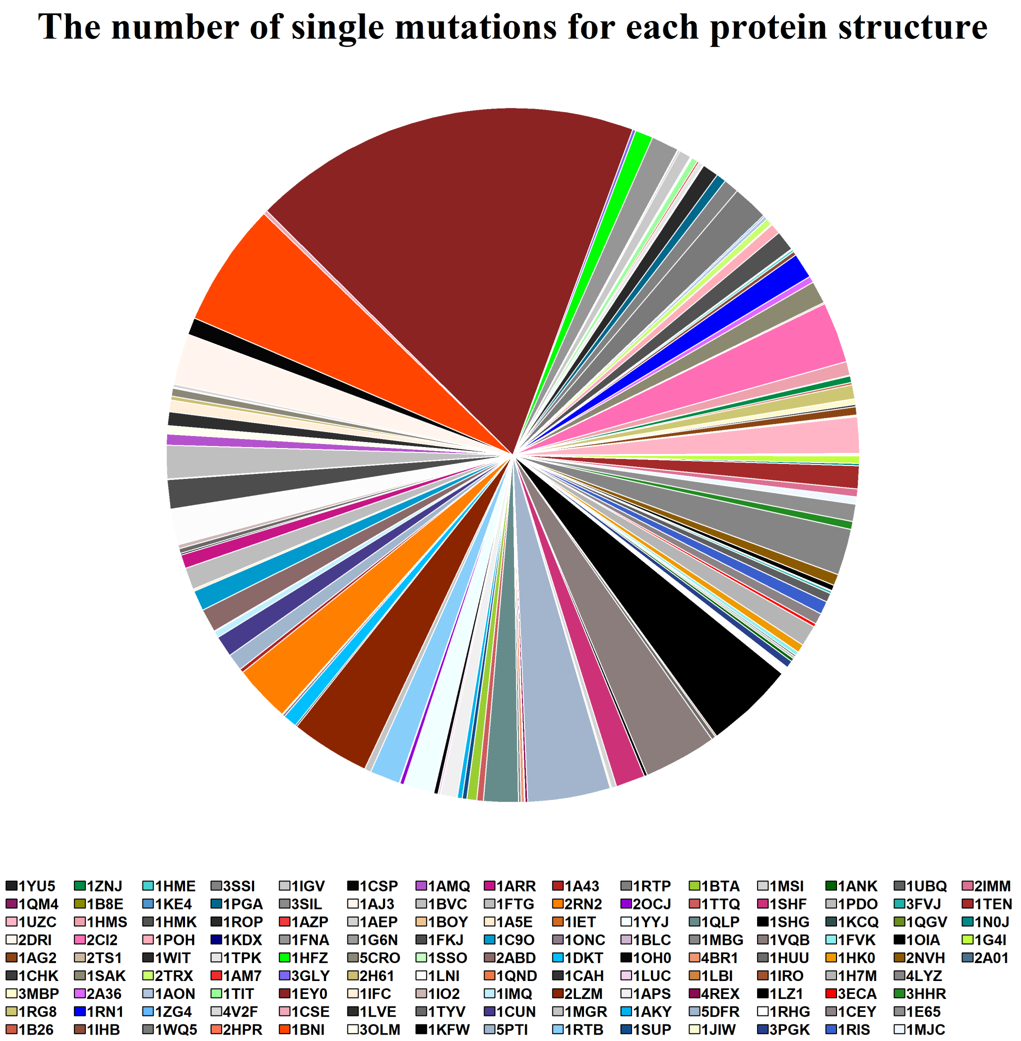


**Figure S1.** The number of mutations for each protein structure in S2648 dataset.

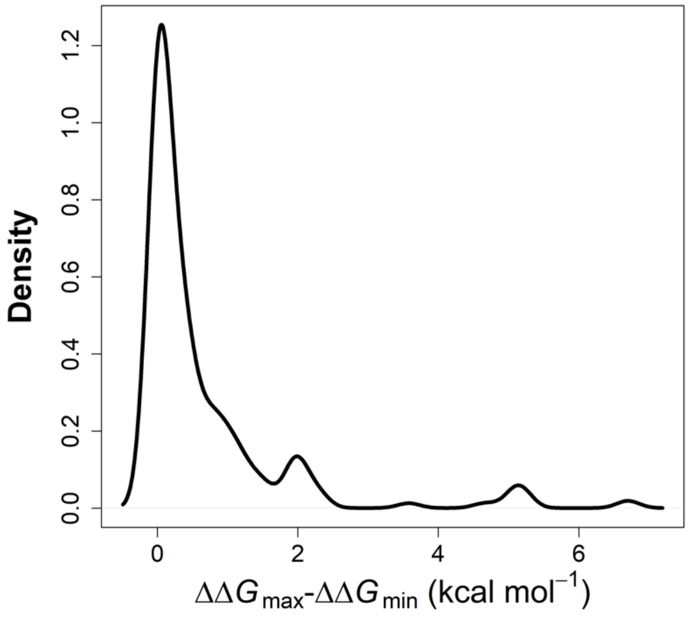


**Figure S2.** The distribution of the differences between maximal and minimal experimentally-determined stability changes ( $\Delta\Delta G_{\max}$ - $\Delta\Delta G_{\min}$ ) for 232 mutations with multiple experimental measurements. The values of $\Delta\Delta G_{\max}$ - $\Delta\Delta G_{\min}$ of 205 mutations are less than 1.0 kcal mol$^{-1}$.

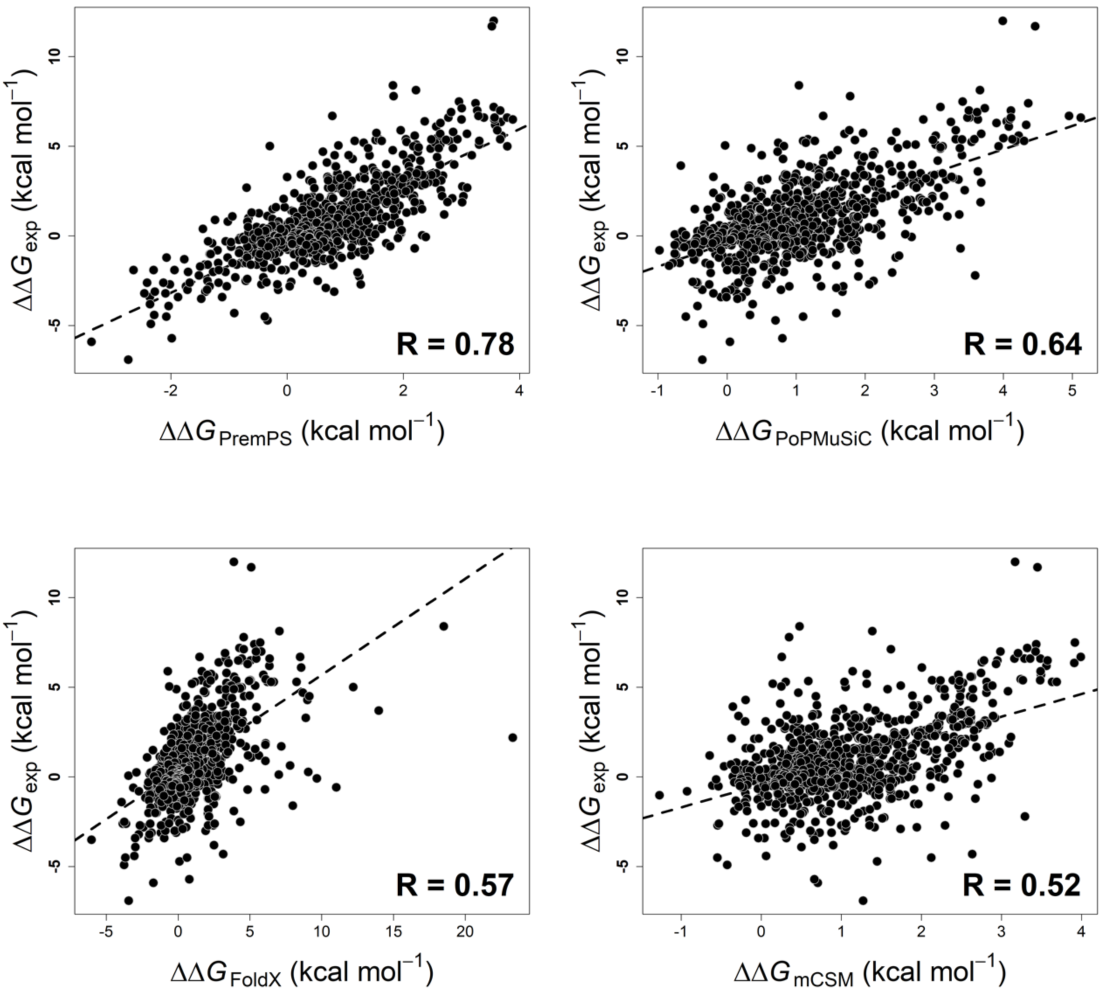


**Figure S3.** Pearson correlation coefficients between experimental and calculated protein stability changes ($\Delta\Delta G$) for PremPS, PoPMuSiC, FoldX and mCSM methods tested on the mutations from S921 dataset, respectively.

**a. Test on S5296**

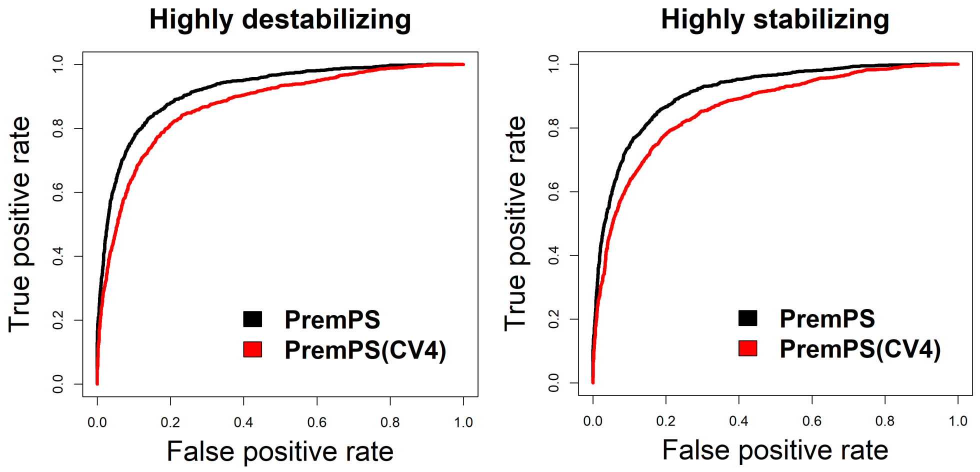


**b. Test on S921**

| Method | Highly destabilizing | | Highly stabilizing | |
|---|---|---|---|---|
| | AUC | MCC | AUC | MCC |
| PremPS | 0.88 | 0.61 | 0.87 | 0.58 |
| PoPMuSiC | 0.81 | 0.49 | 0.67 | 0.21 |
| FoldX | 0.80 | 0.49 | 0.72 | 0.35 |
| mCSM | 0.75 | 0.44 | 0.61 | 0.14 |

**c.**

| Category | | Definition | # of mutations | |
|---|---|---|---|---|
| | | | S5296 | S921 |
| **Highly destabilizing** | **Positive** | $\Delta\Delta G_{exp}$ (kcal mol$^{-1}$) $\geq 1$ | 1364 | 360 |
| | **Negative** | $\Delta\Delta G_{exp}$ (kcal mol$^{-1}$) $< 1$ | 3932 | 561 |
| **Highly stabilizing** | **Positive** | $\Delta\Delta G_{exp}$ (kcal mol$^{-1}$) $\leq -1$ | 1364 | 109 |
| | **Negative** | $\Delta\Delta G_{exp}$ (kcal mol$^{-1}$) $> -1$ | 3932 | 812 |

**Figure S4.** ROC analysis for predicting highly destabilizing and stabilizing mutations. (a) ROC curves for PremPS trained and tested on S5296 and applying leave-one-protein-out validation (CV4) on S5296. (b) AUC and MCC values for different methods tested on S921. The difference of AUC between PremPS and other methods is significant (p-value << 0.01, DeLong test). Maximum Matthews correlation coefficient is calculated for each method. (c) The definition and the number of mutations for making ROC curves.

**a.**

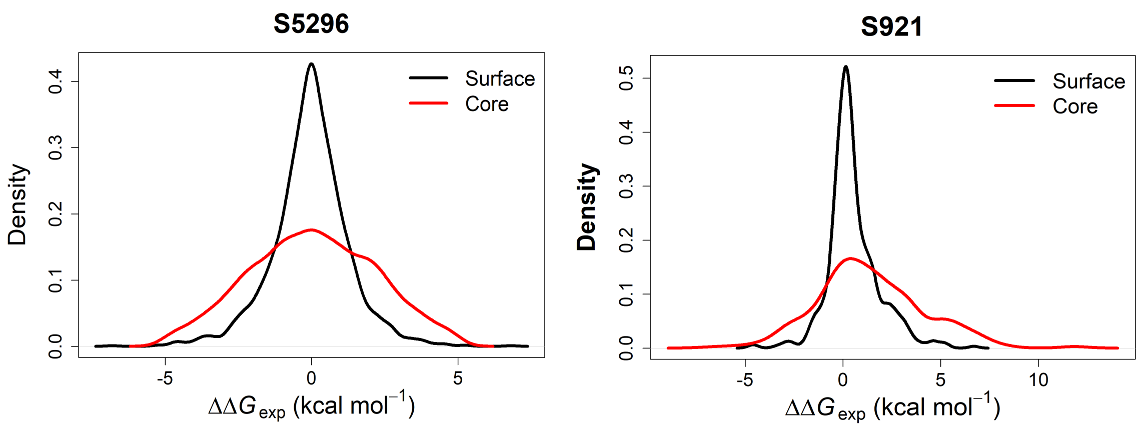


**b.**

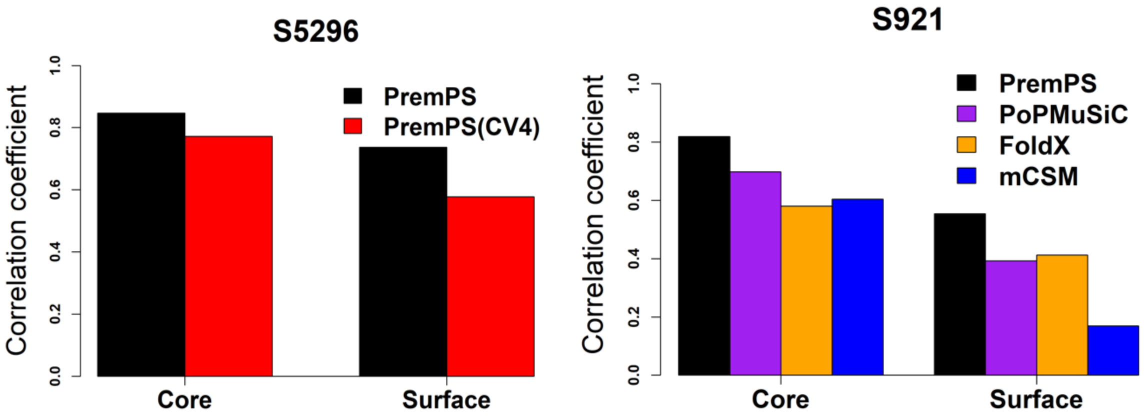


**c.**

| **Category** | **# of mutations** | |
|---|---|---|
| | **S5296** | **S921** |
| **Core** | 2706 | 430 |
| **Surface** | 2590 | 491 |

**Figure S5.** (a) Distribution of experimental values of protein stability changes for mutations located in core and surface respectively. (b) Pearson correlation coefficients between experimental and calculated $\Delta\Delta G$ values. The difference in R between PremPS and other methods is significant (p-value < 0.01, Hittner2003). (c) The number of core and surface mutations in S5296 and S921 datasets, respectively.

**a.**

| Dataset | All mutations | | Non-neutral | | Neutral | |
|---|---|---|---|---|---|---|
| | # of mutations | # of proteins (# of mapped structures) | # of mutations | # of proteins (# of mapped structures) | # of mutations | # of proteins (# of mapped structures) |
| F5276 | 5276 | 58 | 1139 | 46 | 4137 | 46 |
| F2196 | 2196 | 50 (1554) | 1037 | 38 (1388) | 1159 | 39 (1201) |

**b.**

| Method | AUC-ROC | AUC-PR | MCC |
|---|---|---|---|
| PremPS (maximum) | 0.88 | 0.83 | 0.60 |
| PremPS (average) | 0.73 | 0.69 | 0.37 |
| VEST4 | 0.82 | 0.80 | 0.53 |
| B-Score | 0.77 | 0.80 | 0.51 |
| CHASMplus | 0.79 | 0.80 | 0.48 |
| VEST3 | 0.77 | 0.75 | 0.46 |
| REVEL | 0.76 | 0.75 | 0.46 |
| MutationAssessor | 0.76 | 0.74 | 0.46 |
| MutationTaster | 0.76 | 0.50 | 0.45 |
| MutPred | 0.77 | 0.76 | 0.44 |
| CHASM | 0.76 | 0.71 | 0.43 |
| PROVEAN | 0.78 | 0.73 | 0.43 |
| phyloP | 0.74 | 0.67 | 0.42 |
| Eigen | 0.77 | 0.72 | 0.41 |
| PolyPhen2-HVAR | 0.76 | 0.64 | 0.41 |
| LRT | 0.73 | 0.46 | 0.41 |
| PolyPhen2-HDIV | 0.75 | 0.53 | 0.39 |
| CADD | 0.74 | 0.67 | 0.38 |
| MetaLR | 0.66 | 0.67 | 0.37 |
| FatHMM | 0.68 | 0.65 | 0.34 |
| CanDrAplus | 0.63 | 0.56 | 0.30 |
| SiPhy | 0.68 | 0.60 | 0.30 |
| SIFT | 0.68 | 0.44 | 0.30 |
| GERP++ | 0.67 | 0.59 | 0.29 |
| GenoCanyon | 0.64 | 0.57 | 0.27 |
| DANN | 0.65 | 0.59 | 0.25 |
| phastCons | 0.62 | 0.31 | 0.25 |
| M-CAP | 0.63 | 0.62 | 0.24 |
| MetaSVM | 0.54 | 0.45 | 0.17 |
| fitCons | 0.56 | 0.53 | 0.14 |

**c.**

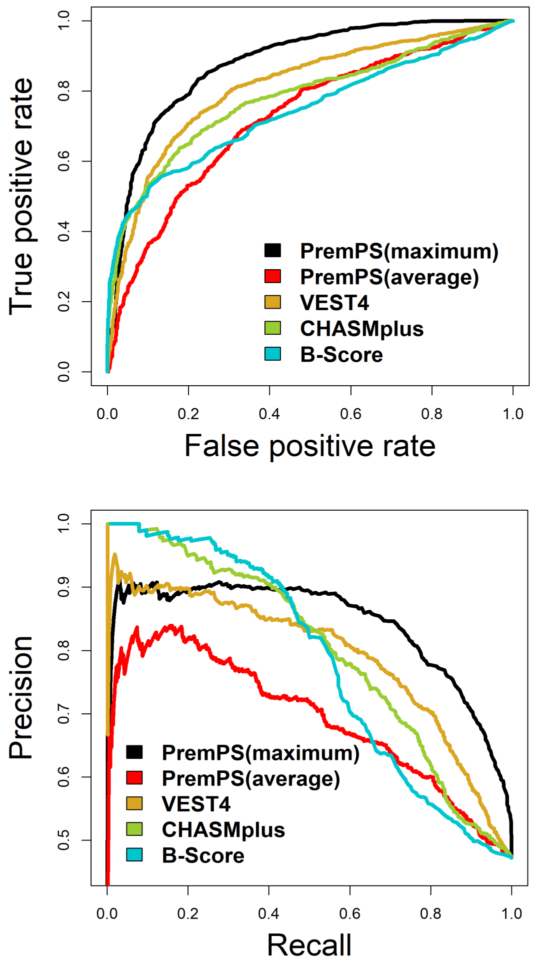

**Figure S6.** Comparison between PremPS and other 28 computational methods in distinguishing non-neutral from neutral mutations. (a) F5276: the previously proposed experimental dataset including 1139 non-neutral/deleterious and 4137 neutral/benign mutations (1); F2196: a dataset is composed of those mutations in F5276 that could be mapped to the corresponding protein 3D structures (one mutation/protein could be mapped to several PDB structures). (b) The performance of PremPS and other 28 computational methods in distinguishing non-neutral from neutral mutations using the dataset of F2196. PremPS (maximum): the maximum predicted absolute value of $\Delta\Delta G$ was used for each non-neutral mutation and minimum predicted absolute value was used for each neutral mutation; PremPS (average): the average value of protein stability changes calculated using all mapped structures was used. Since some methods failed to calculate the scores for some mutations, 1856 mutations for which the predictions are available for all methods were used for calculating AUC-ROC, AUC-PR and maximum MCC values. The methods are ranked with respect to the MCC. (c) ROC and Precision-Recall curves for PremPS, VEST4, CHASMplus and B-Score methods tested on F2196 dataset.

**a.**

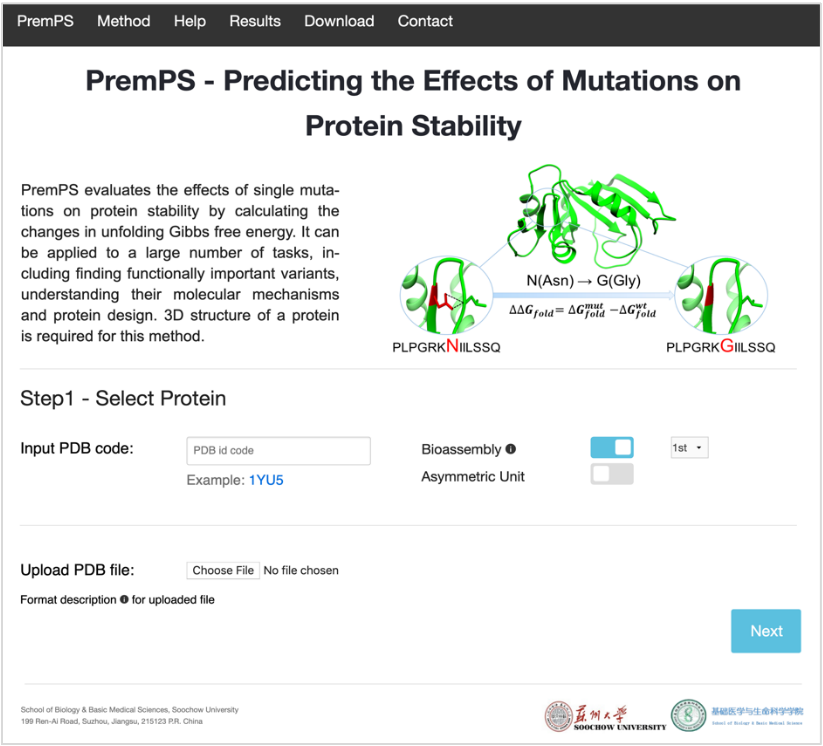
PremPS
Method
Help
Results
Download
Contact
PremPS - Predicting the Effects of Mutations on Protein Stability
PremPS evaluates the effects of single mutations on protein stability by calculating the changes in unfolding Gibbs free energy. It can be applied to a large number of tasks, including finding functionally important variants, understanding their molecular mechanisms and protein design. 3D structure of a protein is required for this method.
N(Asn) → G(Gly)
$\Delta\Delta G_{fold} = \Delta G_{fold}^{mut} - \Delta G_{fold}^{wt}$
PLPGRKNIILSSQ
PLPGRKGIILSSQ
Step1 - Select Protein
Input PDB code:
PDB id code
Example: 1YU5
Bioassembly
Asymmetric Unit
1st
Upload PDB file:
Choose File
No file chosen
Format description for uploaded file
Next
School of Biology & Basic Medical Sciences, Soochow University
199 Ren-Ai Road, Suzhou, Jiangsu, 215123 P.R. China
SOOCHOW UNIVERSITY

**b.**

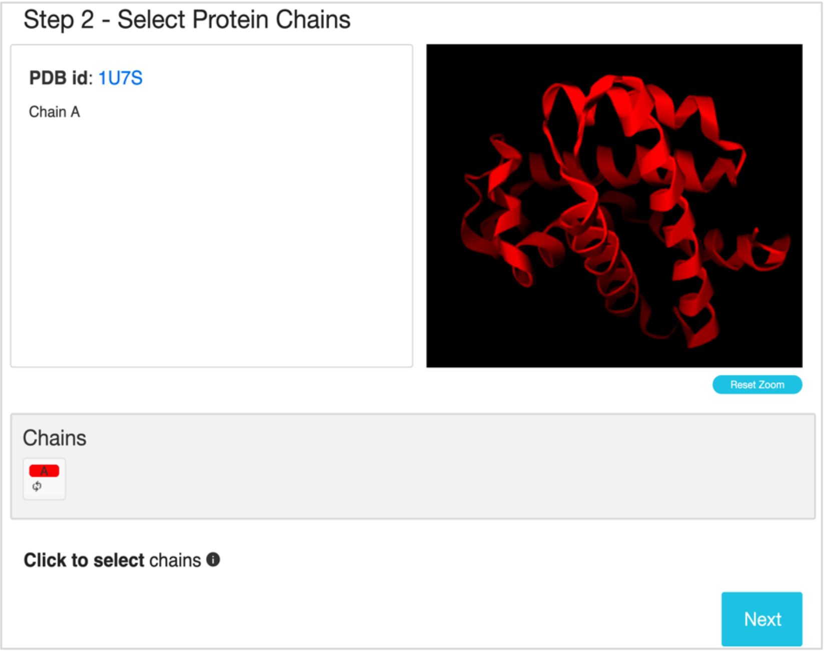
Step 2 - Select Protein Chains
PDB id: 1U7S
Chain A
Reset Zoom
Chains
A
Click to select chains
Next

**c.**

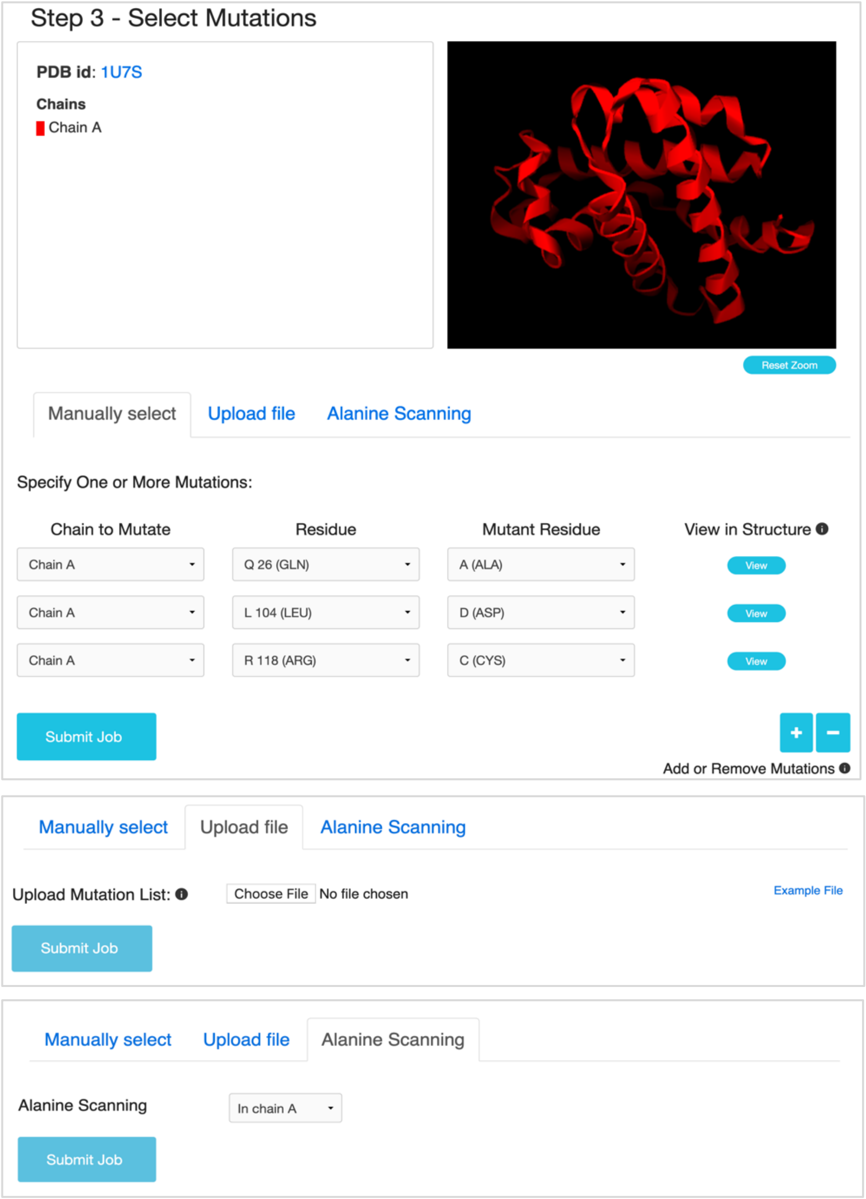


**Figure S7.** (a) The entry page of PremPS server. (b) The second step for selecting protein chains. (c) The third step for selecting mutations and three options are provided: "Specify One or More Mutations Manually", "Upload Mutation List" and "Alanine Scanning for Each Chain".

**a.**

#### Job id: 2019121211341159727269857

- Summary

| PDB ID | Chains | Number of mutations | Start time (EST) | Process time | Results |
|---|---|---|---|---|---|
| 1U7S | A | 3 | 2019-12-12 06:41 | 3 min | Download |

- Results

| # | Mutated Chain | Mutation | ΔΔG | Location | Structure |
|---|---|---|---|---|---|
| 1 | A | Q26A | **-0.64** | COR | Explore |
| 2 | A | L104D | **1.82** | COR | Explore |
| 3 | A | R118C | **0.27** | SUR | Explore |

**b.**

### Non-covalent Interactions Viewer

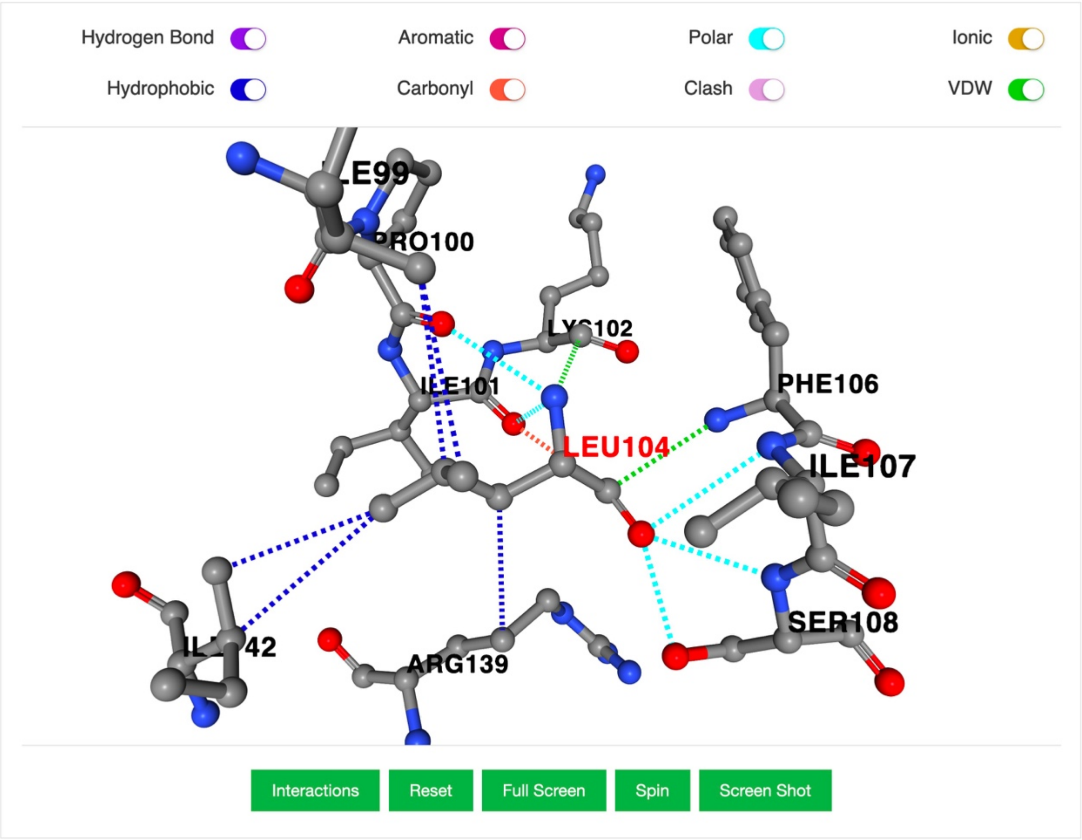


Description for each non-covalent interaction is shown in here.

**Figure S8.** (a) The final results. The contribution of each feature is provided in the download file (b) An interactive 3D viewer showing the non-covalent interactions between the mutated site of Leu104 in the protein myoglobin (PDB ID: 1U7S) and its adjacent residues, generated by Arpeggio.

**Table S1. Experimental datasets used for training and testing.**

a. The number of mutations and protein structures in each dataset.

| Dataset | All mutations | | | $\Delta\Delta G_{exp} \geq 0$ | | $\Delta\Delta G_{exp} < 0$ | |
|---|---|---|---|---|---|---|---|
| | **# of mutations** | **# of proteins** | **# of structures** | **# of mutations** | **# of structures** | **# of mutations** | **# of structures** |
| **S2648** | 2648 | 129 | 131 | 2080 | 119 | 568 | 93 |
| **S5296** | 5296 | 129 | 131 | 2682 | 131 | 2614 | 130 |
| **S350** | 350 | 66 | 67 | 260 | 57 | 90 | 35 |
| **S605** | 605 | 55 | 58 | 467 | 54 | 138 | 34 |
| **S1925** | 1925 | 53 | 55 | 1373 | 48 | 552 | 42 |
| **$S^{sym}$** | 684 | 14 | 357 | 350 | 103 | 334 | 262 |
| **S134** | 134 | 1 | 6 | 98 | 6 | 36 | 6 |
| **p53** | 42 | 1 | 1 | 31 | 1 | 11 | 1 |
| **S921** | 921 | 54 | 195 | 634 | 82 | 287 | 145 |

One mutation/protein could be mapped to several PDB structures.

b. The number of mutations (protein structures) in the training dataset of S5296 that overlaps with each test set.

| Dataset | S350 | S605 | S1925 | $S^{sym}$ | S134 | *p*53 |
|---|---|---|---|---|---|---|
| **S5296** | 350 (67) | 277 (40) | 903 (45) | 402 (14) | 41 (1) | 5 (1) |

**Table S2. The importance of each category of features for PremPS model.** IncNodePurity is used for describing the importance which is the total decrease in node impurities from splitting on the variable, averaged over all trees.

| Feature | Importance |
|---|---|
| $\Delta CS$ | 4909 |
| $\Delta OMH$ | 2502 |
| $PSSM$ | 2398 |
| $SASA_{pro}$ and $SASA_{sol}$ | 2605 |
| $P_{FWY}$, $P_{RKDE}$ and $P_L$ | 2544 |
| $N_{Hydro}$ and $N_{Charg}$ | 1228 |

**Table S3. The performance using Random Forest (RF), Support Vector Machine (SVM) and eXtreme Gradient Boosting (XGBoost) algorithms to build PremPS model and tested on S5296 and S921.**

| Algorithm | Test set | Method | All mutations | | | $\Delta\Delta G_{exp} \geq 0$ | | $\Delta\Delta G_{exp} < 0$ | |
|---|---|---|---|---|---|---|---|---|---|
| | | | R | RMSE | Slope | R | RMSE | R | RMSE |
| **RF** | S5296 | PremPS (CV4) | 0.73 | 1.23 | 1.04 | 0.54 | 1.20 | 0.50 | 1.25 |
| | S921 | PremPS | 0.78 | 1.48 | 1.52 | 0.72 | 1.54 | 0.60 | 1.33 |
| **SVM** | S5296 | PremPS (CV4) | 0.70* | 1.27 | 0.97 | 0.52* | 1.26 | 0.49 | 1.29 |
| | S921 | PremPS | 0.73* | 1.53 | 1.34 | 0.67* | 1.59 | 0.55* | 1.39 |
| **XGBoost** | S5296 | PremPS (CV4) | 0.71* | 1.25 | 0.99 | 0.53* | 1.22 | 0.50 | 1.28 |
| | S921 | PremPS | 0.77 | 1.45 | 1.40 | 0.71 | 1.51 | 0.59 | 1.32 |

R: Pearson correlation coefficient between experimental and predicted $\Delta\Delta G$ values. RMSE (kcal mol$^{-1}$): root-mean square error. Slope: the slope of the regression line between experimental and predicted $\Delta\Delta G$ values. All presented values of correlation coefficients are statistically significantly different from zero (p-value << 0.01). *p-value < 0.01 compared to Random Forest (Hittner2003 test).

**Table S4. The performance for PremPS training and applying four types of cross-validation (CV1-CV4) on S5296 dataset.**

| Method | All mutations | | | $\Delta\Delta G_{exp} \geq 0$ | | $\Delta\Delta G_{exp} < 0$ | | Core | | Surface | |
|---|---|---|---|---|---|---|---|---|---|---|---|
| | R | RMSE | Slope | R | RMSE | R | RMSE | R | RMSE | R | RMSE |
| PremPS | 0.82 | 1.03 | 1.08 | 0.67 | 1.01 | 0.63 | 1.04 | 0.85 | 1.15 | 0.74 | 0.88 |
| PremPS (CV1) | 0.81 | 1.05 | 1.07 | 0.65 | 1.04 | 0.62 | 1.06 | 0.84 | 1.17 | 0.72 | 0.90 |
| PremPS (CV2) | 0.80 | 1.09 | 1.08 | 0.63 | 1.07 | 0.60 | 1.10 | 0.83 | 1.21 | 0.70 | 0.93 |
| PremPS (CV3) | 0.74 | 1.21 | 1.10 | 0.56 | 1.20 | 0.54 | 1.22 | 0.78 | 1.34 | 0.62 | 1.03 |
| PremPS (CV4) | 0.73 | 1.23 | 1.04 | 0.54 | 1.20 | 0.50 | 1.25 | 0.77 | 1.37 | 0.58 | 1.06 |

Core/Surface: the mutations occurred in the protein core/surface.

**Table S5. Comparative performance of different methods on datasets of S350 (a), S605 (b), S1925 (c), $S^{sym}$ (d), S134 (e) and p53 (f).** PremPS: the model was trained on S5296; $PremPS^T$: the model was retrained after removing each test set and the corresponding reverse mutations from S5296.

**a.** All methods were retrained after removing S350 from their training sets.

| Method | Number of predictions | 350 | | 309 | | 87 | |
|---|---|---|---|---|---|---|---|
| | | R | RMSE | R | RMSE | R | RMSE |
| **$PremPS^T$** | **350** | **0.72** | **1.09** | **0.74** | **1.09** | **0.81** | **1.52** |
| mCSM | 350 | 0.73# | 1.08 | 0.74# | 1.10 | 0.82# | 1.48 |
| MAESTRO | 350 | 0.70# | 1.13 | 0.69# | 1.17 | 0.76# | 1.67 |
| PoPMuSiC v2.0 | 350 | 0.67# | 1.16 | 0.67# | 1.19 | 0.71# | 1.67 |
| PoPMuSiC v1.0 | 350 | 0.62 | 1.24 | 0.63 | 1.25 | 0.70# | 1.66 |
| SDM2 | 350 | 0.61 | 1.29 | 0.61 | 1.32 | 0.69# | 1.71 |
| SDM | 350 | 0.52 | 1.80 | 0.53 | 1.81 | 0.63 | 2.11 |
| Dmutant | 350 | 0.48 | 1.81 | 0.47 | 1.87 | 0.57 | 2.31 |
| AUTOMUTE | 315 | 0.46 | 1.43 | 0.45 | 1.46 | 0.45 | 1.99 |
| CUPSAT | 346 | 0.37 | 1.91 | 0.35 | 1.96 | 0.50 | 2.14 |
| Eris | 334 | 0.35 | 4.12 | 0.34 | 4.28 | 0.49 | 3.91 |
| I-Mutant v2.0 | 346 | 0.29 | 1.65 | 0.27 | 1.69 | 0.27 | 2.39 |

The values of R and RMSE for other methods except for PremPS were taken from (2-5).

350 mutations were tested using every method, while some methods failed to compute the $\Delta\Delta G$ for some mutations, so the predicted $\Delta\Delta G$ values were set to zero when counted these mutations. 309 mutations for which the $\Delta\Delta G$ values are available for all methods, and among them 87 mutations whose experimental $|\Delta\Delta G|$ are $\geq$ 2 kcal $mol^{-1}$. The difference in R between PremPS and other methods is significant except #p-value > 0.05 (Fisher1925 test).

**b.** S605 is the training dataset of Meta-predictor. The R and RMSE reported by Meta-predictor are mean values across 1000 tests. Namely, Meta-predictor randomly chose 50% mutations from S605 as training and used the remaining mutations for testing; the procedure was repeated 1000 times. Other methods except $PremPS^T$ were applied to the S605 directly without removing S605 from their training sets.

| Method | R | RMSE |
|---|---|---|
| **PremPS** | **0.80** | **1.34** |
| $PremPS^T$ | 0.70 | 1.51 |
| Meta-predictor | 0.73 | 1.29 |
| PoPMuSiC v2.0 | 0.68 | 1.32 |
| DFire | 0.64 | 1.84 |
| CUPSAT | 0.55 | 1.77 |
| FoldX | 0.54 | 1.78 |
| Rosetta | 0.54 | 2.34 |
| MultiMutate | 0.54 | 2.34 |
| EGAD | 0.52 | 1.61 |
| I-Mutant v3.0 | 0.51 | 1.52 |
| MUPRO | 0.49 | 1.52 |
| SDM | 0.46 | 1.96 |
| Hunter | 0.32 | 1.89 |

The values of R and RMSE for other methods except for PremPS were taken from (6).
The difference in R between PremPS and other methods is significant (p-value < 0.01, Fisher1925 test).

**c.** S1925 is the training dataset of AUTOMUTE. The R and RMSE for all methods are the results of 20-fold cross-validation on S1925.

| Method | R | RMSE |
|---|---|---|
| **$PremPS^{20}$** | **0.87** | **0.90** |
| mCSM | 0.82 | 1.00 |
| AUTOMUTE (REPTree) | 0.79 | 1.10 |
| AUTOMUTE (SVMreg) | 0.76 | 1.20 |
| I-Mutant v2.0 | 0.71 | 1.30 |

The values of R and RMSE for other methods except for PremPS were taken from (3).
The difference in R between $PremPS^{20}$ and other methods is significant (p-value < 0.01, Fisher1925 test).

**d.** All methods except $PremPS^T$ were applied to the dataset of $S^{sym}$ directly.

| Method | Forward mutations | | Reverse mutations | | |
|---|---|---|---|---|---|
| | R | RMSE | R | RMSE | $R_{FR}$ |
| **PremPS** | **0.81** | **0.96** | **0.73** | **1.12** | **-0.93** |
| $PremPS^T$ | 0.64 | 1.21 | 0.56 | 1.30 | -0.90 |
| MUPRO | 0.79# | 0.94 | 0.07 | 2.51 | -0.02 |
| STRUM | 0.75 | 1.05 | -0.15 | 2.51 | 0.34 |
| AUTOMUTE | 0.73 | 1.07 | -0.01 | 2.61 | -0.06 |
| NeEMO | 0.72 | 1.08 | 0.02 | 2.35 | 0.09 |
| iStable | 0.72 | 1.10 | -0.08 | 2.28 | -0.05 |
| Rosetta | 0.69 | 2.31 | 0.43 | 2.61 | -0.41 |
| FoldX | 0.63 | 1.56 | 0.39 | 2.13 | -0.38 |
| PoPMuSiC v2.1 | 0.63 | 1.21 | 0.25 | 2.18 | -0.29 |
| DUET | 0.63 | 1.20 | 0.13 | 2.38 | -0.21 |
| I-Mutant v3.0 | 0.62 | 1.23 | -0.04 | 2.32 | 0.02 |
| mCSM | 0.61 | 1.23 | 0.14 | 2.43 | -0.26 |
| MAESTRO | 0.52 | 1.36 | 0.32 | 2.09 | -0.34 |
| SDM | 0.51 | 1.74 | 0.32 | 2.28 | -0.75 |
| $PoPMuSiC^{sym}$ | 0.48 | 1.58 | 0.48 | 1.62 | -0.77 |
| CUPSAT | 0.39 | 1.71 | 0.05 | 2.88 | -0.54 |

The values of R and RMSE for other methods except for PremPS were taken from (7).
$R_{FR}$ is the Pearson correlation coefficient between predicted $\Delta\Delta G$ values of the forward and reverse mutations. A non-biased prediction should have $R_{FR}$ equal to -1. The difference in R between PremPS and other methods is significant except #p-value > 0.05 (Fisher1925 test).

**e.** All methods except $PremPS^T$ were applied to the dataset of S134 directly.

| Method | 1A6G | 1A6M | 1BZ6 | 1BZP | 1U7S | 2EKT | AVR |
|---|---|---|---|---|---|---|---|
| **PremPS** | **0.71** | **0.72** | **0.73** | **0.73** | **0.75** | **0.73** | **0.73** |
| $PremPS^T$ | 0.65 | 0.65 | 0.65 | 0.65 | 0.65 | 0.64 | 0.65 |
| I-Mutant v2.0 | 0.65# | 0.65# | 0.64# | 0.65# | 0.64# | 0.65# | 0.65# |
| SDM | 0.58# | 0.58 | 0.60# | 0.57 | 0.60 | 0.59 | 0.59 |
| PoPMuSiC v2.1 | 0.54 | 0.55 | 0.56 | 0.57 | 0.55 | 0.55 | 0.56 |
| I-Mutant v3.0 | 0.54 | 0.54 | 0.55 | 0.54 | 0.53 | 0.54 | 0.54 |
| CUPSAT | 0.36 | 0.31 | 0.25 | 0.30 | 0.45 | 0.48 | 0.40 |
| mCSM | 0.35 | 0.39 | 0.40 | 0.44 | 0.47 | 0.44 | 0.44 |

The values of R and RMSE for other methods except for PremPS were taken from (8).
Pearson correlation coefficients between experimental and predicted $\Delta\Delta G$ values for different methods applied on 134 mutations from six high-resolution structures of myoglobin (PDB IDs are shown in the first row). AVR: correlation when using the average $\Delta\Delta G$ values of the six outputs. The difference in R between PremPS and other methods is significant except #p-value > 0.05 (Fisher1925 test).

**f.** All methods except $PremPS^T$ were applied to the dataset of p53 directly.

| Method | R | RMSE |
|---|---|---|
| **PremPS** | **0.73** | **1.41** |
| $PremPS^T$ | 0.71 | 1.49 |
| DUET | 0.68 | 1.39 |
| mCSM | 0.68 | 1.40 |
| PoPMuSiC v2.0 | 0.56 | 1.52 |
| SDM | 0.52 | 1.61 |
| iStable | 0.49 | 1.59 |

The values of R and RMSE for other methods except for PremPS were taken from (3,9).

The difference in R between PremPS and other methods is not significant (p-value > 0.05, Fisher1925 test).

**Table S6. Comparison of methods' performance on the independent test set of S921.** PremPS: PremPS model is trained on S5296 dataset; PremPS$^R$: PremPS model is retrained on the forward mutation dataset of S2648.

| Method | All mutations | | | $\Delta\Delta G_{exp} \geq 0$ | | $\Delta\Delta G_{exp} < 0$ | | Core | | Surface | |
|---|---|---|---|---|---|---|---|---|---|---|---|
| | **R** | **RMSE** | **Slope** | **R** | **RMSE** | **R** | **RMSE** | **R** | **RMSE** | **R** | **RMSE** |
| PremPS | 0.78 | 1.48 | 1.52 | 0.72 | 1.54 | 0.60 | 1.33 | 0.82 | 1.81 | 0.55 | 1.11 |
| PremPS$^R$ | 0.73 | 1.60 | 1.76 | 0.73[a] | 1.51 | 0.29 | 1.79 | 0.80[b] | 2.00 | 0.50 | 1.15 |
| PoPMuSiC | 0.64 | 1.68 | 1.30 | 0.68[c] | 1.48 | - | 2.06 | 0.70 | 2.09 | 0.39 | 1.22 |
| FoldX | 0.57 | 2.06 | 0.54 | 0.56 | 1.99 | 0.22 | 2.21 | 0.58 | 2.58 | 0.41 | 1.45 |
| mCSM | 0.52 | 1.85 | 1.26 | 0.57 | 1.63 | - | 2.25 | 0.60 | 2.28 | 0.17 | 1.36 |

Only correlation coefficients with statistically significantly different from zero (p-value << 0.01, t-test) are shown. The difference in R between PremPS and other methods is significant (p-values < 0.01, Hittner2003) except [a]p-value = 0.27, [b]p-value = 0.02 and [c]p-value = 0.04.
